# Evaluation of Model Fit of Inferred Admixture Proportions

**DOI:** 10.1101/708883

**Authors:** Genís Garcia-Erill, Anders Albrechtsen

## Abstract

Model based methods for genetic clustering of individuals such as those implemented in *structure* or ADMIXTURE allow to infer individual ancestries and study population structure. The underlying model makes several assumptions about the demographic history that shaped the analysed genetic data. One assumption is that all individuals are a result of K homogeneous ancestral populations that are all well represented in the data, while another assumption is that no drift happened after the admixture event. The histories of many real world populations do not conform to that model, and in that case taking the inferred admixture proportions at face value might be misleading. We propose a method to evaluate the fit of admixture models based on estimating the correlation of the residual difference between the true genotypes and the genotypes predicted by the model. When the model assumptions are not violated, the residuals from a pair of individuals are not correlated. In case of a bad fit, individuals with similar demographic histories have a positive correlation of their residuals. Using simulated and real data, we show how the method is able to detect a bad fit of inferred admixture proportions due to using an insufficient number of clusters K or to demographic histories that deviate significantly from the admixture model assumptions, such as admixture from ghost populations, drift after admixture events and non-discrete ancestral populations. We have implemented the method as an open source software that can be applied to both unphased genotypes and next generation sequencing data.

## 1 Introduction

Characterization of population structure is a key step in population genetics. It is essential for understanding the population histories, and it is important to take into account when analysing genetic data, for example in genetic association studies (Lawson et al., 2019). Model-based clustering of individuals is a widely used approach to do so, with several programs available (Alexander, Novembre, & Lange 2009, Pritchard, Stephens, & Donnelly, 2000; Skotte, Korneliussen, & Albrechtsen, 2013; Tang, Peng, Wang, & Risch, 2005). Each software has different inference procedures and other variations, but they all share a common likelihood model, which we will refer to as *admixture model*. Under this model, each allele is an independent sample from a binomial distribution, with probability equal to an *individual allele frequency* specific to each individual and loci. From a pre-defined number of clusters *K* individual frequencies are characterized by the individual admixture proportions *Q* and the ancestral population allele frequencies *F*.

The admixture model implies several assumptions about the demographic history behind the analysed individuals. First, it assumes that all individuals derive from *K* homogeneous ancestral populations, all of them in Hardy-Weinberg equilibrium (HWE) and well represented in the data set. Moreover, individuals are assumed to be unrelated and no linkage disequilibrium (LD) among loci is expected, although there are some extensions of the original model that assume and make use of LD between loci (Falush, Stephens, & Pritchard, 2003). In cases of admixture, i.e. when individuals are assigned to more than one cluster, the ancestral and present population frequencies are assumed to be the same, corresponding to a recent admixture event where the effect of genetic drift is negligible. At the same time, the source ancestral population of the two alleles of an individual at the same loci are treated as independent, which does not account for the presence of very recent hybridization. All these model assumptions will not be a good approximation to the real demographic history of many populations, which calls for a model selection procedure to check whether the model is a good fit to the data it is applied to.

Most of the model selection work regarding the admixture model has been focused in choosing the optimal value of *K* to model the data. Several model selection procedures have been proposed (Alexander & Lange, 2011; Evanno, Regnaut, & Goudet, 2005; Pritchard et al., 2000; Raj, Stephens, & Pritchard, 2014; Verity & Nichols, 2016; Wang, 2019), but none has lead to an unambiguous general solution to the problem (Janes et al., 2017; Novembre, 2016). Moreover, many population histories do not conform to the model. In that case there does not really exist anything similar to a correct *K*, and applying the admixture model can lead to misleading conclusions. A recent paper (Lawson, van Dorp, & Falush, 2018) showed and discussed several cases of problematic admixture analyses, and introduced badMIXTURE, a tool to evaluate the model fit of admixture proportions. This method requires applying CHROMOPAINTER (Lawson, Hellenthal, Myers, & Falush, 2012) and comparing its results with what has been obtained from the admixture model. CHROMOPAINTER is based on patterns of phased haplotype sharing, and therefore requires having phased genotype data, the presence of LD in the data and the existence of linkage maps for the analysed organism. badMIXTURE can still be applied to unlinked loci, but its resolution decreases sensibly in that case, and it requires a certain amount of markers to detect some signal (Lawson et al., 2018). Its requirements are therefore out of reach to many contexts in which the admixture model is commonly used, such as in the study of non-model organisms and in general any study making use of cost-efficient approaches like low-depth next generation sequencing (NGS).

We propose an alternative method to asses the admixture model fit, explicitly based on evaluating to what extent the estimated individual allele frequencies are accurately capturing the true individuals genetic ancestry, and allowing in most cases for an assessment of the model fit at the individual level. It can easily be applied to any situation where the admixture model is used, including when working with low-depth NGS data, where the analysis need to be done from genotype likelihoods to avoid the bias introduced by calling genotypes (Skotte et al., 2013). After introducing the method, we show on simulated and real data how it can detect a bad admixture model fit due to using an insufficient number of ancestral clusters *K* or to demographic histories that deviate from the model assumptions, as well as detect the presence of related individuals in the sample.

## 2 Methods

### 2.1 Correlation of residuals as a measure of admixture model fit

For *N* individuals, *M* sites and *K* ancestral populations, we have an *N × M* genotype matrix *G*, and we have estimated with the admixture model an *N × K* admixture proportions matrix 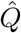 and an *M × K* ancestral frequencies matrix 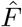. We assume for convenience that genotypes indicate the number of minor alleles individual *i* carries at site *j*; so *g_ij_* ∈ {0,1,2}. Under the admixture model, each genotype *g_ij_* is a sample from a binomial distribution with parameters *n* = 2 and 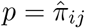, where 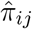is the estimated individual allele frequency of individual *i* at site *j*, and comes from an *M × N* matrix 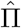 with its elements given by

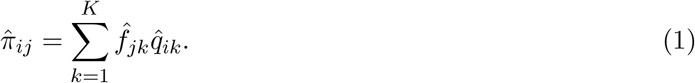

Here 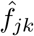 and 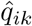 are entries in the 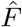 and 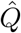 matrices, indicating the allele frequency for site *j* in ancestral population *k* and the admixture proportion of individual *i* from ancestral population *k*, respectively. Let 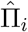 be a vector containing the estimated individual allele frequencies of all sites for individual *i*. We treat each of the true genotypes as a realization from a binomial distribution with probability given by unknown true individual frequencies Π, and *n* = 2. The expected true genotype will be 𝔼[*g_ij_*] = 2*π_ij_*. As an evaluation of the admixture model fit, we aim to test, for each individual *i*, if 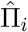 are unbiased estimates of the true Π_*i*_.

We define the predicted genotypes 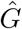, an *N ×M* matrix containing the expectation of the genotypes conditional on the estimated individual frequencies

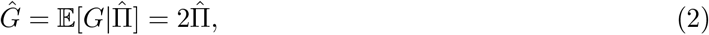

and *R*, another *N ×M* residual matrix containing the difference between the true and the predicted genotypes

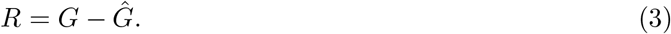

*R_*i*_* is a vector containing the residuals of all *M* sites from individual *i*. If there is a good admixture model fit, i.e. 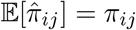, the expected residual 𝔼[*r_*ij*_*] = 0 for every site. But with a bad model fit, the expected estimated frequency will deviate from the true frequency by some quantity 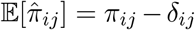; and this will be present in the expected residuals

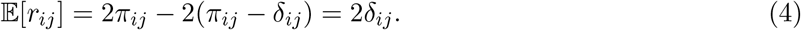

This deviation will be specific to every individual and site, and the mean residual of an individual across sites can be zero even when there is a bad fit. However, we can detect cases of a bad fit by considering that individuals from the same population, with similar genetic backgrounds that are not accurately described by the admixture model, will tend to share the same specific deviations at all sites. As a measure of admixture model fit, we use the correlation of residuals between pairs of individuals, e.g. between individuals 1 and 2,

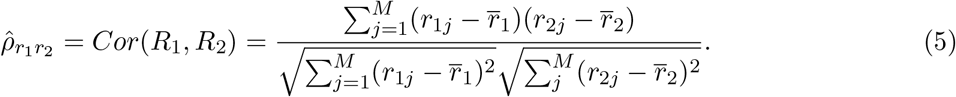

If individual 1 and 2 are from the same population, under a bad fit the sharing of a systematic error in their individual frequencies will result in a positive correlation of their residuals. With a good model fit, which we define as having unbiased estimated individual allele frequencies, the residuals would only be a result of the binomial variance, and they would be independent between any pair of individuals.

### 2.2 Correlation of residuals as a measure of relatedness

A pair of related individuals, that share one or more recent common ancestors, will tend to share more alleles identical by descent (IBD) than a pair randomly drawn from the same population. Related individuals have therefore correlated genotypes, which results in a positive correlation of residuals between that pair even if their ancestry is accurately described by the individual allele frequencies, i.e. even when there is a good admixture model fit.

To show how the correlation of residuals detects relatedness, we can compare it with the population structure-robust kinship estimator 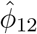 used in REAP (Thornton et al., 2012) and PC-relate (Conomos, Reiner, Weir, & Thornton, 2016). This estimator accounts for population structure by conditioning the gentoype correlation on the individual allele frequencies. The kinship coefficient then is equal to half the genotype correlation; for individuals 1 and 2 it is 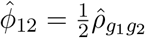 (Thornton et al., 2012). If we assume that we have estimated accurate admixture proportions for both individuals 1 and 2, we have that their expected residual is zero and we can disregard the mean residual in the correlation formula (5). In that case the correlation of residuals is equal to the genotype correlation, 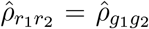, and therefore for individuals with a good model fit the correlation of residuals would be twice their kinship coefficient.

### 2.3 Frequency correction

Although the residuals between unrelated individuals with a good model fit should be independent, this is not the case when using the initial ancestral frequencies as estimated with the admixture model. The correlation of residuals in this case contains a bias that arrives from the estimation of the ancestral frequencies 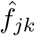. The ancestral frequencies 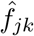 are estimated using the genotypes of all individuals with some admixture proportion from population *k*, which introduces a bias in the correlation of the residuals between individuals that share ancestry from one or more populations. For example if individuals 1 and 2 are non-admixed individuals assigned to the same population *k*, the expected correlation under a good model fit is

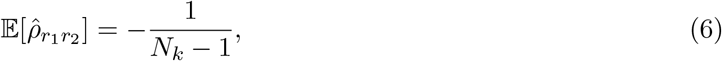

where *N_*k*_* is the number of individuals assigned to ancestral population *k*, assuming that all individuals with ancestry from that population are not admixed (See Appendix S1 for details).

We can make the residuals between individuals 1 and 2 independent under a good model fit, by removing the contribution of individual 1 from the frequency used to calculate individual 2’s residuals. Assuming for simplicity that the genome of every individual comes from a single ancestral population *k*, for a site *j*

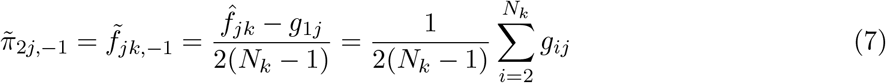

In this case, when 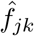 is an unbiased estimate of the true frequency *f_*jk*_*, individual’s 1 initial residuals 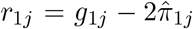 and individual’s 2 corrected residuals 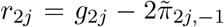 are independent and we expect a zero correlation of their residuals (proof in Appendix S2).

We apply this correction extending it to allow for the presence of admixed individuals, whose contribution to each of the ancestral population allele frequencies is given by the probability that their alleles come from that population. This probability depends on the individual’s genotype and admixture proportions and on the ancestral population allele frequencies. The estimation of the probability requires several iterations of an expectation maximization (EM) algorithm, which introduces higher computational requirements to the method. See appendix Appendix S3 for a detailed description on how the frequency correction is done with admixed individuals.

### 2.4 Method with NGS data

Calling genotypes when using low or medium depth NGS data can introduce several biases in population genetic analyses, including the admixture model (Nielsen, Paul, Albrechtsen, & Song, 2011; Skotte et al., 2013). These biases can be avoided by working directly from the genotype likelihoods. For this reason we implemented a version of the method in a genotype likelihoods framework. Genotype likelihoods give the probability of observing the sequencing data *X* given that the true genotype is *g_*ij*_*; *P* (*X_*ij*_|G_*ij*_* = *g_*ij*_*), and they can be estimated from NGS data with software such as GATK (McKenna et al., 2010), SAMtools (Li, 2011) or ANGSD (Korneliussen, Albrechtsen, & Nielsen, 2014). From the genotype likelihoods, we can obtain the probability of each genotype given the data

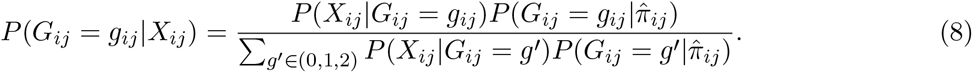

We use the individual frequencies obtained from the admixture results as prior, and assume Hardy-Weinberg equilibrium

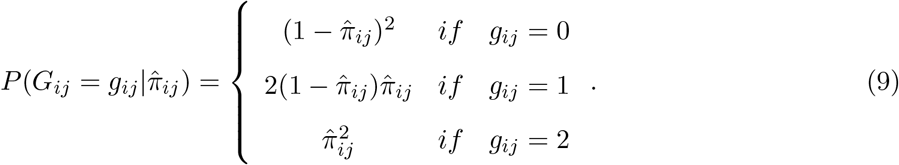

Considering all three possible genotypes, we calculate the expected genotype given the sequencing data

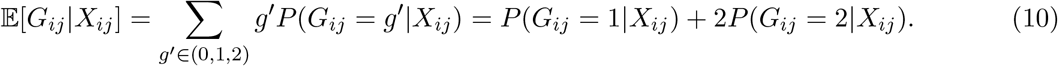

This expected genotype is used instead of the true known genotype in the calculation of the residuals

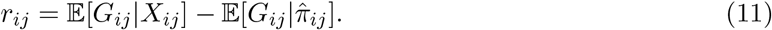

### 2.5 Simulations

#### 2.5.1 Genotypes

To test the method, we simulated genotype data for two different scenarios. The two scenarios were generated from allele frequency data of the Human Genome Diversity Project (HGDP) (Cann et al., 2002). In Scenario 1 we used an European (French), an East Asian (Han Chinese) and an African (Yoruba) populations, and furthermore we created an admixed population with admixture proportions (*Q_Fr_*, *Q_Han_*, *Q_Yor_*) = (0.7, 0.3, 0). In scenario 2 we included also the French, Han and Yoruba popula-tions, and an admixed population that in this case has ancestry from a Native American population (Zapotec), with (*Q_Fr_*, *Q_Han_*, *Q_Yor_*, *Q_Zap_*) = (0.7, 0, 0, 0.3). For each scenario we simulated genotypes for 80 individuals in total from around 0.5 million sites. In both of them there were 20 individuals from the French, the Han and the Yoruba populations, and 20 from the admixed population. The genotypes were generated by sampling from a binomial distribution using the individual frequencies as given by (1), which in non-admixed individuals reduce to the population frequencies. All sites were in HWE and without LD between them. A minimum allele frequency (MAF) filter was applied using PLINK v.1.07 (Purcell et al., 2007); around 0.4 million sites were left in each scenario. Admixture pro-portions and ancestral population frequencies were estimated for both scenarios using ADMIXTURE (Alexander et al., 2009) assuming *K* = 3.

#### 2.5.2 Genotype likelihoods

For each of these genotype datasets, we also simulated low-coverage NGS data by sampling reads with a mean depth of 3*X*, assuming a Poisson distribution. Sequencing errors were included by having a probability *e* = 0.01 of sampling the wrong base. Genotype likelihoods were then calculated from the sampled reads. A more detailed description of the simulation protocol used can be found in (Meisner & Albrechtsen, 2018). Admixture proportions and ancestral allele frequencies were estimated from the genotype likelihoods using NGSadmix (Skotte et al., 2013) with *K* = 3 and default settings.

### 2.6 Badmixture simulations

We obtained the freely available data of the three simulated scenarios (Bottleneck, Ghost Admixture and Recent Admixture) used in (Lawson et al., 2018) as genotypes in PLINK format. The simulations comprise a total of 13 populations that aim at mimicking human population history, originally performed as part of a study on the origin of the Ethiopian Ari populations (van Dorp et al., 2015). Each of the three scenarios differ in the relationships between 4 of these populations, in which we and the previous studies focused and that we label Pop1, Pop2, Pop3 and Pop4. The whole dataset contains a total of 795 individuals in the scenarios labelled as Bottleneck and Ghost Admixture, and 785 in the Recent Admixture. Regarding the four populations of interest, the Bottleneck and Ghost Admixture scenario contain 15 individuals in Pop1, 25 in Pop2, 100 in Pop3 and 25 in Pop4, and in Recent Admixture there are 35 individuals in Pop1, 25 in Pop2, 70 in Pop3 and 25 in Pop4. We used PLINK v.1.07 to filter variants with MAF below 0.05 and pruned markers in LD by removing one of each pair of sites with an *r*^2^ above 0.1 within windows of 100 kb; about 40, 000 sites were left after filtering in each scenario. For each of the scenarios, we ran ADMIXTURE in the dataset including all 13 populations assuming *K* = 11, and calculated the correlation of residuals between individuals from Pop1, Pop2, Pop3 and Pop4.

### 2.7 1000 Genomes Data

#### 2.7.1 Genotypes

We also tested the method on real human data from the 1000 Genomes Project Consortium (1000G) (Auton et al., 2015). We used 435 individuals from 5 populations: 108 from Yoruba in Ibadan, Nigeria (YRI), 61 from African Ancestry in Southwest US (ASW), 99 from Utah residents with Northern and Western European ancestry (CEU), 63 from Mexican Ancestry in Los Angeles, California (MXL) and 103 from Han Chinese in Beijing, China (CHB). We used autosomal sites from the Human Origins panel (Lazaridis et al., 2014), with 407, 441 sites left after filtering sites with minor allele frequency (MAF) below 0.05. We ran ADMIXTURE with *K* = 3 and *K* = 4, and calculated the correlation of residuals in each case. We inferred relatedness between all pairs of individuals in the dataset using relateAdmix (Moltke & Albrechtsen, 2014), that is based on inferring the fraction of sites in which a pair of individuals have 1 (*k*_1_) or 2 (*k*_2_) alleles identical by descent (IBD) while accounting for population structure and admixture using the estimated admixture proportions and ancestral allele frequencies. Kinship coefficients can then be calculated as 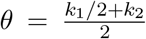. We used the admixture parameters estimated with *K* = 4 to control for the effect of admixture in relatedness inference.

#### 2.7.2 Genotype likelihoods

To test the NGS version of the method, we used low-coverage sequencing data from the same 1000G populations, individuals and sites used for the genotype version. We calculated genotype likelihoods using ANGSD (Korneliussen et al., 2014), applying the genotype likelihood model described in McKenna et al., 2010, and filtering bases with sequence quality below 30 and reads with mapping quality below 20, excluding sites without data in more than 50 individuals, with a SNP significance threshold of 1*e*^−6^ and excluding variants with MAF below 0.05. A total of 408, 240 SNPs were left after filtering. The mean sequencing depth per individual ranged from 2.6*X* to 15.8*X*, with a median of 6.8*X*. We ran NGSadmix assuming *K* = 3 and *K* = 4, and calculated the correlation of residuals between individuals for each model.

## 3 Results

### 3.1 Simulations

We simulated two similar scenarios, that differ in the demographic history of the admixed populations. In Scenario 1 there is admixture between the French and Han populations, while in Scenario 2 it is the unsampled Zapotec population that admixes with the French. This second scenario is therefore a case of admixture from a ghost population which should result in a bad model fit and correlated residuals.

Both scenarios result in nearly identical estimated admixture proportions. The individuals from the admixed populations are inferred as having ancestry from the French and the Han populations (Figures 1A and 2A). This results are accurately capturing the simulated genetic background for all individuals in Scenario 1, and we obtain uncorrelated residuals between them (Figure 1C, visualized as shown in Figure 1B). In Scenario 2 the inferred admixture proportions are not an accurate description of the admixed individuals’ demographic history. The individual frequencies are here calculated using the allele frequencies from the Han ancestral population instead of the Zapotec from which these individuals were simulated to have ancestry for. This leads to positively correlated residuals between the admixed individuals (Figure 2B).

**Figure 1:**
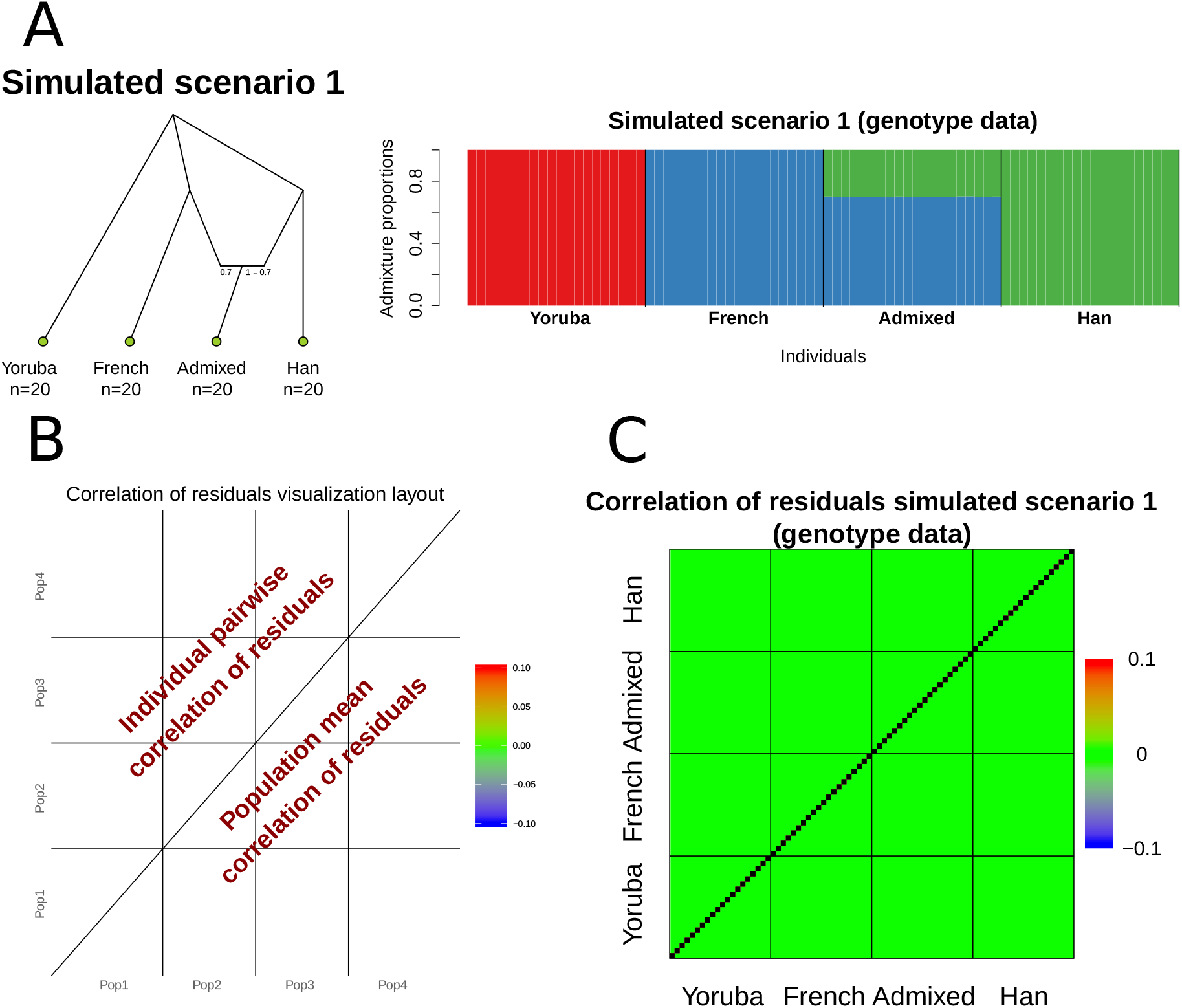

Admixture model results and evaluation of simulated scenario 1. A. Population tree of the simulated demographic history in Scenario 1 and inferred admixture proportions assuming *K* = 3 from genotype data using ADMIXTURE. B. Diagram with the placement of data in the heatmap used for visualizing the correlation of residuals C. Evaluation of model fit of the admixture results as the correlation of residuals: residuals are independent between all individuals, indicating a good admixture model fit.

**Figure 2:**
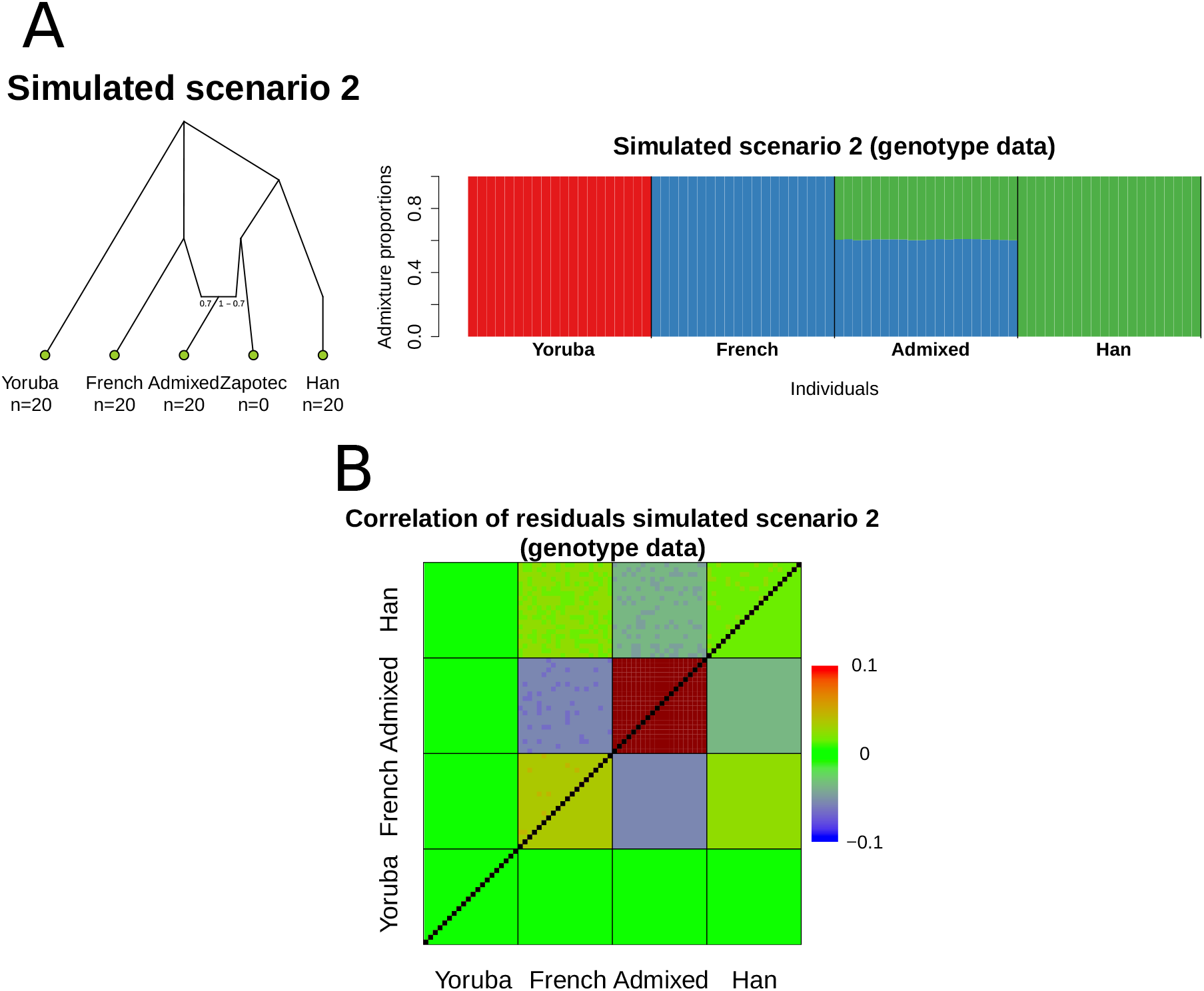

Admixture model results and evaluation of simulated scenario 2. A. Population tree of the simulated demographic history in Scenario 2 and inferred admixture proportions assuming *K* = 3 from genotype data using ADMIXTURE. B. Evaluation of model fit of the admixture results as the correlation of residuals; a positive correlation within the admixed population, and to a lesser extend in the two source populations, indicate a bad fit of the inferred admixture proportions. Correlation values below or above the color scale are plotted as dark blue and dark red, respectively

We also applied the method to genotype likelihoods obtained from simulated NGS data from the same datasets with a mean depth of 3*X*, obtaining similar results (Supplementary Figures S1 and S2).

### 3.2 BadMIXTURE simulations

The simulations used in the BadMIXTURE paper comprise three scenarios with different demographic histories. Yet, all three scenarios result in nearly identical estimated admixture proportions when the admixture model is applied, with Pop2 being modelled as a mixture of Pop1, Pop3 and Pop4 (Figure 3A). After running the admixture model in these simulated datasets, we calculated the correlation of residuals for each scenario.

**Figure 3:**
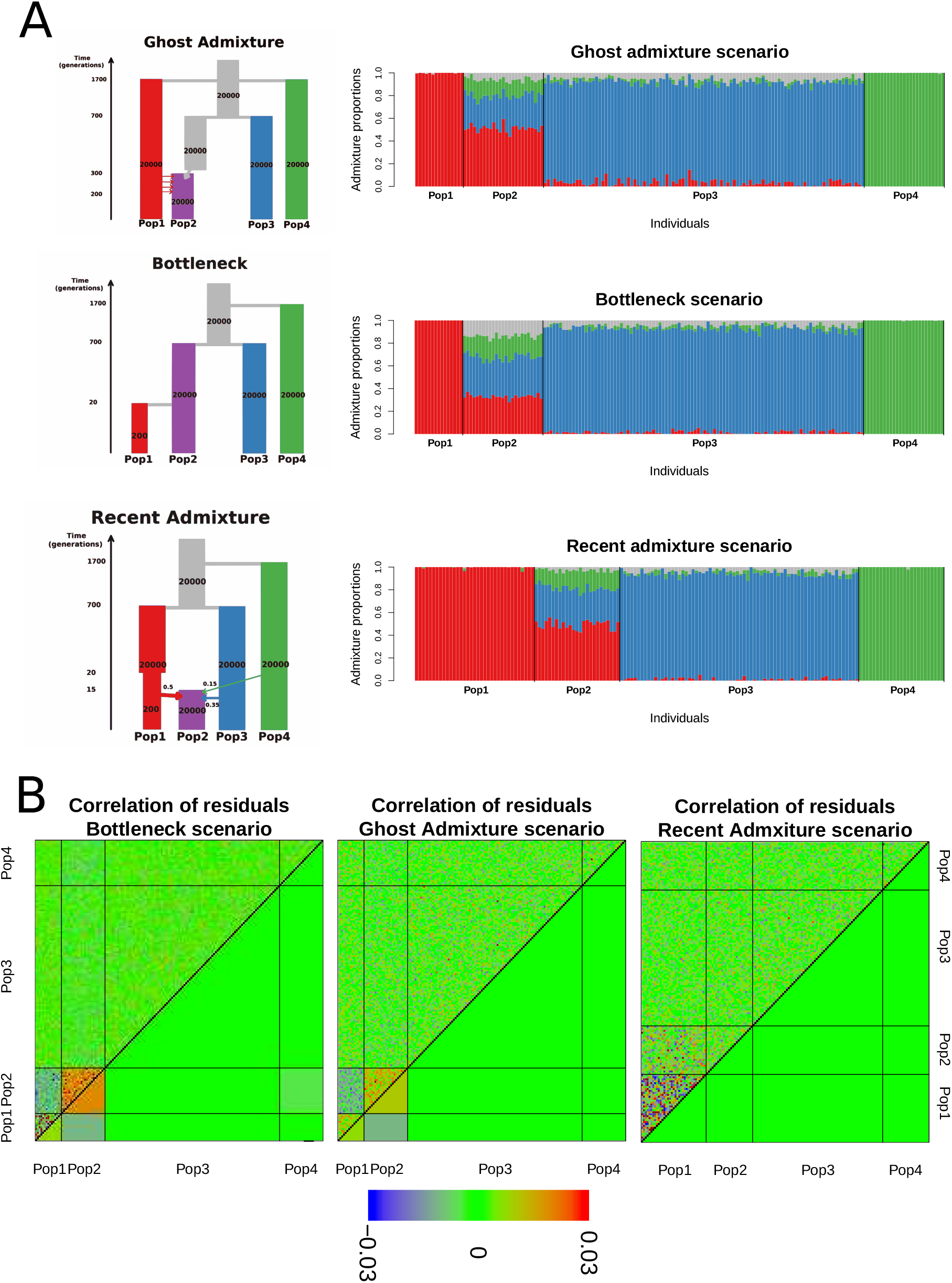

Admixtuer model results and evaluation of the three simulated scenarios from Lawson, van Dorp, and Falush, 2018. A. Demographic models of each of the three scenarios, including only the four populations of interest (time and population sizes are not scaled), with the corresponding inferred admixture proportions assuming *K* = 11, colouring only the three relevant ancestral populations, and pooling together the rest as grey. B. Evaluation of the estimated admixture proportions for each scenario as the correlation of residuals. Correlation values below or above the color scale are plotted as dark blue and dark red, respectively.

In the Bottleneck scenario, a recent bottleneck in Pop1 results in Pop2 being modelled as admixed despite the absence of any gene flow. We can detect this bad model fit as a positive correlation of residuals between individuals from Pop2 (Figure 3B). In the Ghost admixture scenario, Pop2 has received admixture from an unsampled population. Again a positive correlation of residuals shows that the admixture results are not an accurate representation of its demographic history (Figure 3B). In the Recent admixture scenario, Pop2 is the result of an admixture event between Pop1, Pop2 and Pop3 in proportions similar to the inferred ones. In this case we find a zero mean correlation of residuals between individuals from Pop2, and from the rest of populations (Figure 3B).

### 3.3 1000 Genomes

We applied the admixture model to a subset of 5 populations (YRI, ASW, CEU, MXL and CHB, see methods) from the 1000 Genomes project, assuming 3 and 4 ancestral clusters. In the case with *K* = 3, the inferred clusters correspond to African, European and East Asian ancestral populations. The Native American ancestry in the MXL population is modelled using the East Asian ancestry as a proxy, which results in a positive correlation of the residuals between individuals from that populations, with magnitude varying depending on the amount of Native American ancestry the individuals have (Figure 4A 4B). Moreover, there are a few individuals from the ASW population with Native American ancestry, who have also a positive correlation with MXL individuals. When doing the analysis with *K* = 4, a cluster corresponding to the Native American ancestry is added. In this case, individuals with Native American ancestry have uncorrelated residuals (Figure 4B). In none of the population there is a visible mean correlation when using a normal scale. However, when increasing the resolution we find a weak signal in the ASW population (Supplementary Figure S3), suggesting that the YRI and CEU populations are not a perfect proxy for the ancestry of African Americans. We ran the analyses using both genotype and NGS data, inferring in both cases similar admixture proportions and correlation of residuals (Supplementary Figure S4).

**Figure 4:**
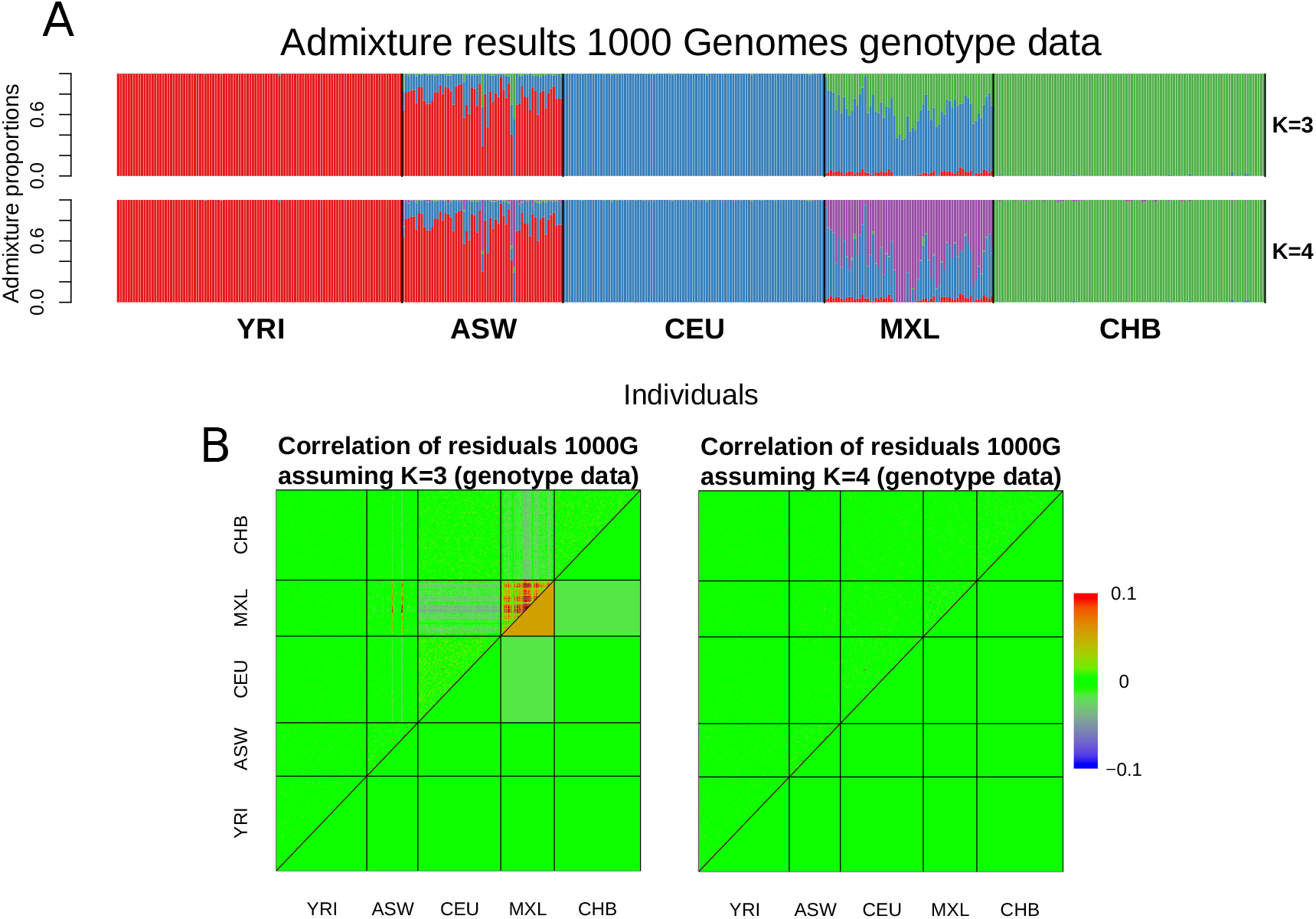

Admixture model results and evaluation of real human data from the 1000G projects. A. Admixture proportions of the genotype data from the 1000 Genomes dataset, assuming *K* = 3 and *K* = 4. B. Evaluation of the inferred admixture proportions as correlation of residuals when assuming *K* = 3 or *K* = 4, showing that four ancestral populations are needed to accurately model the ancestry of individuals from the MXL population. Correlation values below or above the color scale are plotted as dark blue and dark red, respectively. Population abbreviations: YRI Yoruba in Ibadan, Nigeria; ASW African Ancestry in Southwest US; CEU Utah residents with Northern and Western European ancestry; MXL Mexican Ancestry in Los Angeles, California; CHB Han Chinese in Beijing, China.

There are a several individuals in the 1000G analysis that show a high correlation of their residuals with the same magnitude at both *K* = 3 and *K* = 4. The majority of these cases, and also the ones with the strongest signal, are from the ASW population (Figure 4B). We inferred relatedness using relateAdmix, and found five pairs of individuals from the ASW population with a kinship coefficient of 0.25 and one pair with a kinship coefficient of 0.12. These six pairs were the same that had the highest correlation of residuals, and their correlation of residuals is roughly twice the kinship coefficient (Figure 5).

**Figure 5:**
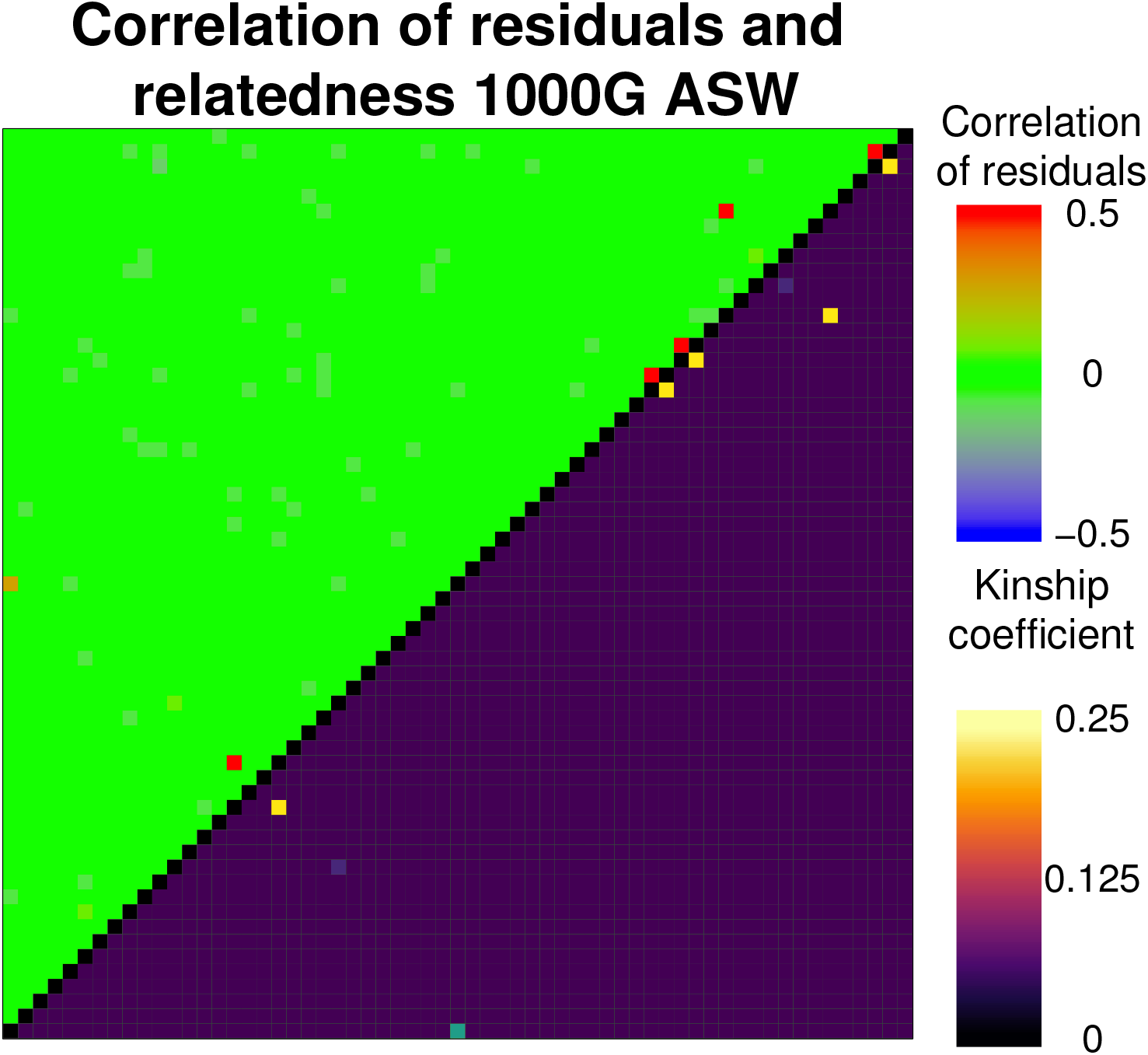

Correlation of residuals between the ASW individuals from the 1000 Genomes dataset with admixture proportions inferred assuming *K* = 4 (above diagonal) and kinship coefficients between the same individuals estimated with relateAdmix (below diagonal). The pairs with highest correlation show also the highest kinship coefficient.

### 3.4 Implementation

The method presented here has been implemented as evalAdmix in C++, and can be run multithreaded. The time complexity is 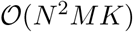, It is quadratic with respect to the number of individuals since it has to estimate ancestral allele frequencies *N* times in contrast which ADMIXTURE which is linear, 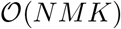. However, it is still fast enough to run on thousands of samples (Supplementary Figure S5). The implementation is freely available at https://github.com/GenisGE/evalAdmix.

## 4 Discussion

We have introduced a method to evaluate the model fit of admixture proportions, and we have shown how it can detect different violations of the admixture model assumptions, such as assuming a too low number of ancestral clusters, drift between recently diverged populations, admixture from ghost populations and the presence of related individuals in the sample. Moreover, the correlation of residuals as a measure of admixture model fit allows for a simple interpretation at the individual level, as it is based on evaluating whether the individual allele frequencies estimated with the admixture model are an accurate description of the individuals genetic ancestry. The magnitude of the correlation will in general indicate how divergent is the individual’s ancestry as described by the model from the true individual’s ancestry. It could potentially be interpreted as in a recently proposed framework where *F_ST_* is defined between individual-specific sub-populations using individual allele frequencies (Ochoa & Storey, 2019). In case of a bad model fit, the correlation of residuals will depend on the individuals’ admixture proportions and on the divergence between the respective estimated ancestral populations and an idealized true population. The latter will furthermore depend on the sample size included from different populations. All this complicates deriving an expression in terms of *F_ST_* that could allow the correlation of residuals to be a quantitative measure of the divergence between the estimated ancestry and the true ancestry.

Above what threshold can it be considered that there is a bad model fit is a question that needs addressing. However, we do not think it possible to give a general binary answer to that question. Real data will never have a perfect fit to the admixture model assumptions, so the question becomes when is the model fit good enough. Our analyses with the 1000G data provides a relevant example. While with *K* = 3 it is clear that we are missing an ancestral cluster to model the Native American ancestry, at *K* = 4 we can still detect a certain correlation in the mean within the ASW population, probably reflecting that there is more diversity in the African and European admixture sources of the African Americans than what is captured using only the YRI and CEU populations as sources (Tishkoff et al., 2009). However, it is a very subtle correlation and we need to reduce the scale of the visualization to a level in which the signal of individual correlations is dominated by noise. For many purposes, we could consider that this signal is negligible enough so that the results with four ancestral clusters can be seen as a good approximation.

Increasing the number of ancestral clusters *K* means using more parameters in the calculation of the individual allele frequencies, which will always lead to a decrease in the total mean correlation of residuals. What the correlation of residuals can provide is a lower bound to the number of ancestral populations needed to model the data, as the smallest value of *K* from which the resulting correlation is good enough. In contrast, most procedures proposed to select *K*, such as *structure*’s model evidence (Pritchard et al., 2000), ADMIXTURE’s cross-validation error (Alexander & Lange, 2011) or the Δ*K* statistic (Evanno et al., 2005), focus on avoiding over-fitting and consequently give an upper bound to *K*, which can lead to underestimating the amount of population substructure present in the data (Janes et al., 2017). The presence of population substructure within clusters would be detected by the correlation of residuals as long as there are at least two individuals from that sub-population. Uneven sample sizes can bias the admixture model, for example by causing highly diverged populations not to be assigned their own cluster if too few individuals are present in the sample (Puechmaille, 2016). Because in this case the genetic ancestry of individuals from these populations would not be accurately described by the admixture model, the correlation of residuals is able to detect biased analyses caused by uneven sample sizes.

An excessive focus on the choice of *K* can lead to the implicit assumption that the demographic history of the sample can be represented as a recent mixture of discrete ancestral populations. For most real populations, this is not true. However, the admixture proportions 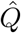 will in any case be used by the model to describe variation in the dataset (Lawson et al., 2018). Therefore, the values in 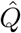 that we call admixture proportions will not always be an estimate of the proportion of each individual’s genome that derives from different source populations. A good fit of the data to the admixture model is needed in order for 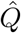 to be taken as representing actual admixture proportions.

We have shown that the correlation of residuals between the six ASW pairs is roughly twice their kinship coefficient, which agrees with the mathematical relationship between the correlation of residuals and the kinship coefficient estimator (Thornton et al., 2012). The five ASW individual pairs with the highest correlation had been previously identified to be parent-offspring, and the sixth pair with a moderate correlation to be aunt/niece (Gazal, Sahbatou, Babron, Genin, & Leutenegger, 2015). The presence of close relatives is a violation of the admixture model assumptions, that can result in biased results by favouring clustering of related individuals (Anderson & Dunham, 2008). Therefore being able to detect related individuals can be useful to detect potential biases in admixture model results. However, it is worth noting that in our 1000G analysis this violation of the model assumption does not change the interpretation of the results. In this case there is enough African and European ancestry in the analysis that the few pairs of related individuals do not create sub clusters or otherwise interfere with the inferred admixture proportions. Related pairs of individuals will have a positive correlation of residuals even if the admixture proportions are accurately capturing the ancestries of both individuals, i.e. even if there is a good model fit at the individual level. Consequently relatedness can potentially be a confounding factor when interpreting the correlation of residuals. There is information in the correlation of residuals that can to some extend aid in distinguishing between positive values due to a bad model fit and to relatedness. In general the signal due to relatedness will only be present in the pair of close relatives, while other individuals from the same population will have uncorrelated residuals, or a lower correlation in case they have a bad fit. However in general definitively distinguishing between relatedness and a bad fit would require using additional information and analyses, such as using methods specific for inferring relatedness.

The method that we propose only requires the input and output used for an admixture analysis, which makes it very simple to integrate them. It can be used to evaluate the results of admixture models applied to SNPs from either genotype and NGS data using *structure* (Pritchard et al., 2000), ADMIXTURE (Alexander et al., 2009), NGSAdmix (Skotte et al., 2013), and in general any implementation of the admixture model that outputs individual admixture proportions and ancestral population allele frequencies. The run-time might in some cases be a problem, since the need to correct the frequencies makes the complexity to increase quadratically with the number of individuals. In most cases it is still reasonably fast, specially if the program can be ran using several CPUs. However, if many thousands of individuals are included in the analysis and the chosen *K* is high it can become a problem. For this cases, we have implemented the option to calculate the correlation only between a selected set of individuals of interest. Additionally, a proportion of the total sites can be specified to be randomly selected, discarding the rest. This decreases linearly the run-time, although if too few sites are left the resolution will also decrease.

We believe the tool we propose can be of great utility in the interpretation of admixture model results. It can provide additional guidance in the persistent problem of choosing an appropriate value of *K* and allows for an assessment of the model fit at the individual level. It is in this sense very similar to badMIXTURE (Lawson et al., 2018), but its lower requirements and ease of use make it an attractive alternative that can be applied in a broader range of situations.

## Supporting information

Frequencies for simulations

1000G individuals IDs

## Acknowledgement

We would like to thank Daniel Falush for the idea for this project which stems from conversations with him at the 2018 Workshop on Population and Speciation Genomics, Cesky Krumlov. We would also like to thank Jonas Meisner for his help in performing the simulations, and Emil Jørsboe for his help in performing the simulations and in solving problems with the implementation. This work was supported by a grant from the Lundbeck Foundation (R215-2015-4174).

## Data Accessibility

The method is freely available at https://github.com/GenisGE/evalAdmix. The frequencies from the HGDP used to sample genotypes for both the genotype and the NGS simulations are included as supporting information. The PLINK files of the simulations from (Lawson et al., 2018) are available at https://people.maths.bris.ac.uk/~madjl/finestructure/badmixture/. The 1000G datasets are available at ftp://ftp.1000genomes.ebi.ac.uk/vol1/ftp/phase3/data/ and ftp://ftp.1000genomes.ebi.ac.uk/vol1/ftp/release/20130502/ for the low coverage sequencing data and the variant callset, respectively. A list with the IDs of the 1000G individuals used in our analyses is included as supporting information.

## Author Contributions

AA and GGE developed the method, GGE implemented the method and performed the analyses with input from AA. GGE wrote the article with input and supervision from AA.

