## Supplementary text and figures for "Evaluation of Model Fit of Inferred Admixture Proportions"

December 5, 2019

#### S1 Appendix 1: Expected correlation using sample frequencies

We assume that we have $N$ diploid genotypes that are sampled from the same panmictic population $k$ with minor allele frequency $f_k$. Genotypes are the minor allele count, so $g \in \{0, 1, 2\}$ and the expected genotype is $\mathbb{E}[g] = 2f$. We use the sampled genotypes to calculate the sample frequency $\hat{f}_k = \frac{1}{2N}\sum_i^N g_i$; with $\mathbb{E}[\hat{f}_k] = f_k$.

The residual $r$ is the difference between the true sampled genotype and the genotype predicted from the estimated sample frequency; so individual 1's residual is

$$r_1 = g_1 - \mathbb{E}[G|\hat{f}_k] = g_1 - \frac{1}{N}\sum_{i=1}^{N} g_i. \tag{1}$$

What we are interested in is the correlation between the residuals of pairs of individuals. Taking individuals 1 and 2, since for any $i$ $\mathbb{E}[r_i] = 0$, we can write the expected correlation $Cor(r_1, r_2) = \hat{\rho}_{r_1 r_2}$ as

$$\mathbb{E}[\hat{\rho}_{r_1 r_2}] = \frac{\mathbb{E}[r_1 r_2]}{\sqrt{\mathbb{E}[r_1 r_1]}\sqrt{\mathbb{E}[r_2 r_2]}} = \frac{\mathbb{E}[(g_1 - \frac{1}{N}\sum_i^N g_i)(g_2 - \frac{1}{N}\sum_i^N g_i)]}{\sqrt{\mathbb{E}[(g_1 - \frac{1}{N}\sum_i^N g_i)(g_1 - \frac{1}{N}\sum_i^N g_i)]}\sqrt{\mathbb{E}[(g_2 - \frac{1}{N}\sum_i^N g_i)(g_2 - \frac{1}{N}\sum_i^N g_i)]}}$$
$$= \frac{\mathbb{E}[g_1 g_2] - \mathbb{E}[g_1 \frac{1}{N}\sum_i^N g_i] - \mathbb{E}[g_2 \frac{1}{N}\sum_i^N g_i] + \mathbb{E}[(\frac{1}{N}\sum_i^N g_i)(\frac{1}{N}\sum_i^N g_i])}{\sqrt{\mathbb{E}[g_1 g_1] - 2\mathbb{E}[g_1 \frac{1}{N}\sum_k^N g_i]\mathbb{E}[(\frac{1}{N}\sum_i^N g_i)(\frac{1}{N}\sum_i^N g_i])}\sqrt{\mathbb{E}[g_2 g_2] - 2\mathbb{E}[g_2 \frac{1}{N}\sum_i^N g_i]\mathbb{E}[(\frac{1}{N}\sum_i^N g_i)(\frac{1}{N}\sum_i^N g_i])}}. \tag{2}$$

We can then expand the expectations of products of sums over genotypes such that

$$\mathbb{E}[g_1 \frac{1}{N}\sum_i^N g_i] = \frac{1}{N}\mathbb{E}[g_1 g_1 + \sum_{j=2}^N g_1 g_j] = \frac{1}{N}(\mathbb{E}[g_1 g_1] + \sum_{j=2}^N \mathbb{E}[g_1 g_j]) \tag{3}$$

and

$$\mathbb{E}[\frac{1}{N}\sum_i^N g_i \frac{1}{N}\sum_i^N g_i] = \frac{1}{N^2}\mathbb{E}[\sum_i^N g_i g_i + \sum_i^N \sum_{j\neq i}^N g_i g_j] = \frac{1}{N^2}(\sum_i^N \mathbb{E}[g_i g_j] + \sum_i^N \sum_{j\neq i}^N \mathbb{E}[g_i g_j]). \tag{4}$$

When assuming that genotypes are drawn from the same population, we can assume them to be IID, and therefore

$$\begin{aligned} \mathbb{E}[g_i] = \mathbb{E}[g_j] = \mathbb{E}[g], \\ \mathbb{E}[g_i g_i] = \mathbb{E}[g_j g_j] = \mathbb{E}[gg], \\ \mathbb{E}[g_i g_j] = \mathbb{E}[g_i]\mathbb{E}[g_j] = \mathbb{E}[g]\mathbb{E}[g], \end{aligned} \tag{5}$$

where $g$ is any genotype of an individual from the same population. When applying this to expansion (3), we can develop it further and denote its value as $d$

$$\frac{1}{N}(\mathbb{E}[g_1 g_1] + \sum_{j=2}^N \mathbb{E}[g_1 g_j]) = \frac{1}{N}(\mathbb{E}[gg] + (N-1)\mathbb{E}[g]\mathbb{E}[g]) = \frac{1}{N}(\mathbb{E}[gg] - \mathbb{E}[g]\mathbb{E}[g]) + \mathbb{E}[g]\mathbb{E}[g] = d. \tag{6}$$

Similarly (4) will have the same value $d$

$$\begin{aligned} \frac{1}{N^2}(\sum_i^N \mathbb{E}[g_i g_i] + \sum_i^N \sum_{j\neq i}^N + \mathbb{E}[g_i g_j]) &= \frac{1}{N^2}(N\mathbb{E}[gg] + N(N-1)\mathbb{E}[g]\mathbb{E}[g]) \\ &= \frac{1}{N}(\mathbb{E}[gg] - \mathbb{E}[g]\mathbb{E}[g]) + \mathbb{E}[g]\mathbb{E}[g] = d. \end{aligned} \tag{7}$$

Going back to the correlation, and initially denoting expectations of products of sums by $d = \frac{1}{N}(\mathbb{E}[gg] - \mathbb{E}[g]\mathbb{E}[g]) + \mathbb{E}[g]\mathbb{E}[g]$:

$$
\begin{aligned}
\mathbb{E}[\hat{\rho}_{r_1 r_2}] = \frac{\mathbb{E}[g]\mathbb{E}[g] - d - d + d}{(\sqrt{\mathbb{E}[gg] - 2d + d})^2} &= \frac{\mathbb{E}[g]\mathbb{E}[g] - d}{\mathbb{E}[gg] - d} \\
&= \frac{\mathbb{E}[g]\mathbb{E}[g] - \frac{1}{N}(\mathbb{E}[gg] - \mathbb{E}[g]\mathbb{E}[g]) - \mathbb{E}[g]\mathbb{E}[g]}{\mathbb{E}[gg] - \frac{1}{N}(\mathbb{E}[gg] - \mathbb{E}[g]\mathbb{E}[g]) - \mathbb{E}[g]\mathbb{E}[g]} \\
&= \frac{-\frac{1}{N}(\mathbb{E}[gg] - \mathbb{E}[g]\mathbb{E}[g])}{\mathbb{E}[gg] - \mathbb{E}[g]\mathbb{E}[g] - \frac{1}{N}(\mathbb{E}[gg] - \mathbb{E}[g]\mathbb{E}[g])} \\
&= -\frac{\frac{1}{N}}{1 - \frac{1}{N}} = -\frac{1}{N-1}.
\end{aligned} \tag{8}
$$

Thus for unadmixed individuals in the same population we expect a negative correlation of the residuals.

#### S2 Appendix 2: Expected correlation with frequency correction

We can make the residuals within one discrete population independent if we remove the contribution of one of the individuals from the frequency used in the calculation of the other individual's residual. So to calculate the correlation between individuals 1 and 2, we can use for individual 1 the sample frequency $\hat{f}_k = \frac{1}{2N}\sum_i^N g_i$, while for individual 2 we use

$$
\tilde{f}_{k,-1} = \frac{1}{2(N-1)}\sum_{i=2}^{N} g_i. \tag{9}
$$

The residuals are then

$$
\begin{aligned}
r_1 &= g_1 - 2\hat{f}_k = g_1 - \frac{1}{N}\sum_i^N g_i, \\
r_{2,-1} &= g_2 - 2\tilde{f}_{k,-1} = g_2 - \frac{1}{N-1}\sum_{i=2}^{N-1} g_i,
\end{aligned} \tag{10}
$$

and the expected correlation $Cor(r_1, r_2) = \hat{\rho}_{r_1 r_2}$ is

$$
\begin{aligned}
\mathbb{E}[\hat{\rho}_{r_1 r_2}] &= \frac{\mathbb{E}[r_1 r_2]}{\sqrt{\mathbb{E}[r_1 r_2]}\sqrt{\mathbb{E}[r_2 r_2]}} = \frac{\mathbb{E}[g_1 - \frac{1}{N}\sum_i^N g_i)(g_2 - \frac{1}{N-1}\sum_{i=2}^N g_i)]}{\sqrt{\mathbb{E}[g_1 - \frac{1}{N}\sum_i^N g_i)(g_1 - \frac{1}{N}\sum_i^N g_i)]}\sqrt{\mathbb{E}[(g_2 - \frac{1}{N-1}\sum_{i=2}^N g_i)(g_2 - \frac{1}{N-1}\sum_{i=2}^N g_i)]}} \\
&= \frac{\mathbb{E}[g_1 g_2] - \mathbb{E}[g_1 \frac{1}{N-1}\sum_{i=2}^N g_i] - \mathbb{E}[g_2 \frac{1}{N}\sum_i^N g_i] + \mathbb{E}[(\frac{1}{N}\sum_i^N g_i)(\frac{1}{N-1}\sum_{i=2}^N g_i)]}{\sqrt{\mathbb{E}[g_1 g_1] - 2\mathbb{E}[g_1 \frac{1}{N}\sum_i^N g_i]\mathbb{E}[(\frac{1}{N}\sum_i^N g_i)(\frac{1}{N}\sum_i^N g_i)]}\sqrt{\mathbb{E}[g_2 g_2] - 2\mathbb{E}[g_2 \frac{1}{N-1}\sum_{i=2}^N g_i]\mathbb{E}[(\frac{1}{N-1}\sum_{i=2}^N g)(\frac{1}{N-1}\sum_{i=2}^N g_i)]}}.
\end{aligned} \tag{11}
$$

To prove that in this case $\mathbb{E}[\hat{\rho}_{r_1 r_2}] = 0$, we only need to show that the expected value of the numerator is 0, since the denominator is the product of two variances, which by definition will have a positive value. There are three expectations of products of sums over genotypes to expand, one of them already given by equation 3, which we denoted as $d$,

$$\mathbb{E}[g_2 \frac{1}{N} \sum_i^N g_i] = \frac{1}{N}(\mathbb{E}[gg] - \mathbb{E}[g]\mathbb{E}[g]) + \mathbb{E}[g]\mathbb{E}[g] = d. \tag{12}$$

Then we have two new expansions

$$\mathbb{E}[g_1 \frac{1}{N-1} \sum_{i=2}^N g_i] = \frac{1}{N-1} \sum_{i=2}^N \mathbb{E}[g_1 g_i] = \frac{1}{N-1}(N-1)\mathbb{E}[g]\mathbb{E}[g] = \mathbb{E}[g]\mathbb{E}[g], \tag{13}$$

and

$$\begin{aligned}\mathbb{E}[(\frac{1}{N} \sum_i^N g_i)(\frac{1}{N-1} \sum_{i=2}^N g_i)] &= \frac{1}{N(N-1)}(\sum_{i=2}^N \mathbb{E}[g_i g_i] + \sum_{i=2}^N \sum_{j=1;j\neq i}^N \mathbb{E}[g_i]\mathbb{E}[g_j]) \\ &= \frac{1}{N}(\mathbb{E}[gg] + (N-1)\mathbb{E}[g]\mathbb{E}[g]) = \frac{1}{N}(\mathbb{E}[gg] - \mathbb{E}[g]\mathbb{E}[g]) + \mathbb{E}[g]\mathbb{E}[g] = d\end{aligned} \tag{14}$$

We denote the non zero denominator as $A$, and by substituting with what we have shown in equations (12), (13) and (14), we find that the expected correlation is

$$\begin{aligned}\mathbb{E}[\hat{\rho}_{r_1 r_2}] = &\frac{\mathbb{E}[g_1 g_2] - \mathbb{E}[g_1 \frac{1}{N-1} \sum_{i=2}^N g_i] - \mathbb{E}[g_2 \frac{1}{N} \sum_i^N g_i] + \mathbb{E}[(\frac{1}{N} \sum_i^N g_i)(\frac{1}{N-1} \sum_{i=2}^N g_i)]}{A} \\ &= \frac{\mathbb{E}[g]\mathbb{E}[g] - d - \mathbb{E}[g]\mathbb{E}[g] + d}{A} = 0.\end{aligned} \tag{15}$$

Thus if we remove the contribution of individual 1 from individual's 2 frequency, their expected residuals will be independent.

#### S3 Appendix 3: Frequency correction with admixed individuals

To extend the frequency correction to the case with admixed individuals, we need to consider that the number of alleles contributed by each individual to the calculation of the ancestral frequency will not be an integer. Consequently, also the sample size $N_k$ used to obtain $f_k$ will not be an integer. Individual's $i$ contribution to the calculation of the allele frequency of site $j$ in population $k$ will depend on the probability that one of its alleles $\mathbb{A} \in \{0, 1\}$ comes from ancestral population $k$. We denote the ancestry of the allele $\mathbb{A}$ of individual $i$ for site $j$ as $A_{ij} = k$ with probability

$$P(A_{ij}=k|\hat{F},\hat{Q},\mathbb{A}=1)=\frac{P(\mathbb{A}=1|A_{ij}=k,\hat{F},\hat{Q})P(A_{ij}=k|\hat{F},\hat{Q})}{\sum_{k'}P(\mathbb{A}=1|A_{ij}=k',\hat{F},\hat{Q})P(A_{ij}=k|\hat{F},\hat{Q})}=\frac{f_{jk}q_{ik}}{\sum_{k'}^{K}f_{jk'}q_{ik'}}, \tag{16}$$

and

$$P(A_{ij}=k|\hat{F},\hat{Q},\mathbb{A}=0)=\frac{P(\mathbb{A}=0|A_{ij}^{1}=k,\hat{F},\hat{Q})P(A_{ij}=k|\hat{F},\hat{Q})}{\sum_{k'}P(\mathbb{A}=0|A_{ij}=k',\hat{F},\hat{Q})P(A_{ij}=k|\hat{F},\hat{Q})}=\frac{(1-f_{jk})q_{ik}}{\sum_{k'}^{K}(1-f_{jk'})q_{ik'}}. \tag{17}$$

The contribution of each individual to the frequency calculation is given by the number of minor and major alleles it carries, given by its genotype, times the probabilities calculated with equations (16) and (17). The estimated ancestral frequencies can be written as

$$\hat{f}_{jk}=\frac{\sum_{i}^{N}P(A_{ij}^{1}=k|\hat{F},\hat{Q},\mathbb{A}=1)g_{ij}}{\sum_{i}^{N}\Big(P(A_{ij}^{1}=k|\hat{F},\hat{Q},\mathbb{A}=1)g_{ij}+P(A_{ij}^{1}=k|\hat{F},\hat{Q},\mathbb{A}=0)(2-g_{ij})\Big)}. \tag{18}$$

Then we can obtain a new frequency estimate without an individual's contribution by excluding that individual from the summation in both the numerator and the denominator. For example the frequency estimate without the contribution of individual 1 is

$$\tilde{f}_{jk,-1}=\frac{\sum_{i=2}^{N}P(A_{ij}=k|\hat{F},\hat{Q},\mathbb{A}=1)g_{ij}}{\sum_{i=2}^{N}\Big(P(A_{ij}=k|\hat{F},\hat{Q},\mathbb{A}=1)g_{ij}+P(A_{ij}=k|\hat{F},\hat{Q},\mathbb{A}=0)(2-g_{ij})\Big)}. \tag{19}$$

Because we used the initial frequencies to obtain the ancestral probabilities with equations (16) and (17), the new estimate will still contain some contribution of individual 1, and therefore the correlation of residuals will still be biased. To obtain an estimate of the unbiased frequencies, we used an expectation maximization (EM) iterative algorithm. At each iteration, the ancestral probabilities calculated with equations (16) and (17) are calculated using the frequency estimated in the previous iteration with equation (19). We used a fixed number of 5 iterations, which is a conservative setting since in all tested scenarios between 3 and 4 iterations were enough to obtain an unbiased correlation, meaning it tended to 0 for unrelated individuals with a good model fit. Using the corrected ancestral frequencies without individual's 1 contribution $\tilde{f}_{jk,-1}$, each entry from individual's 2's individual allele frequencies $\tilde{\Pi}_{2,-1}$ is given by

$$\tilde{\pi}_{2j,-1}=\sum_{k=1}^{K}\tilde{f}_{jk,-1}\hat{q}_{2k} \tag{20}$$

And these frequencies are used to obtain individual's 2 residuals used to calculate the correlation with individual's 1 residuals

$$R_{2,-1} = G_2 - 2\tilde{\Pi}_{2,-1}. \tag{21}$$

The residuals of individual 2 are calculated from the frequencies from the admixture model (**??**). In case of a good admixture model fit,

$$\mathbb{E}[\hat{\rho}_{r_1 r_2}] = \mathbb{E}[Cor(R_1, R_{2,-1})] = 0. \tag{22}$$

### S4 Supplementary Figures

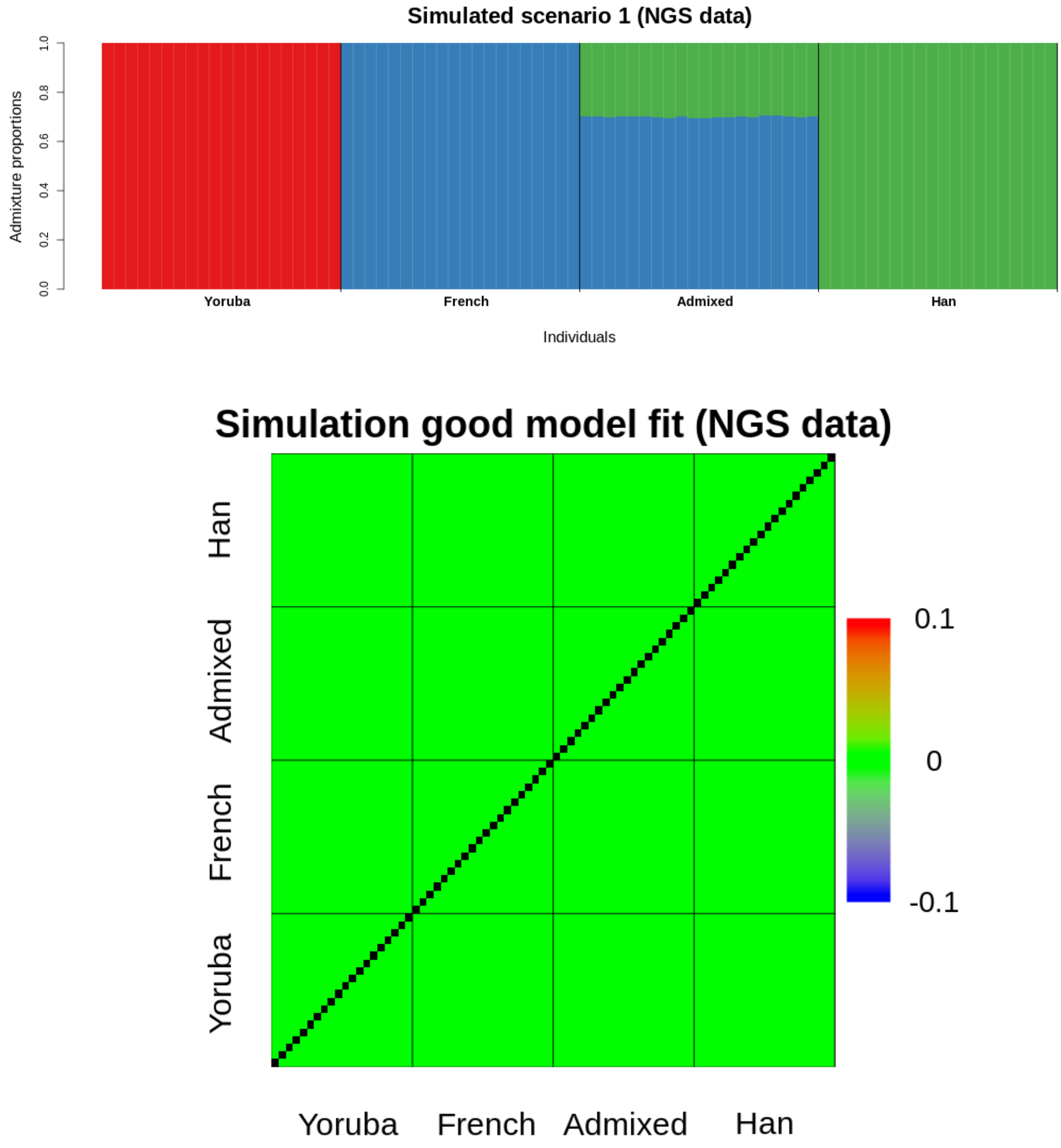


Figure S1: Admixture proportions inferred with NGSadmix assuming $K = 3$ from genotype likelihood data for simulated Scenario 1 (upper panel) and evaluation of model fit as correlation of residuals (bottom panel.

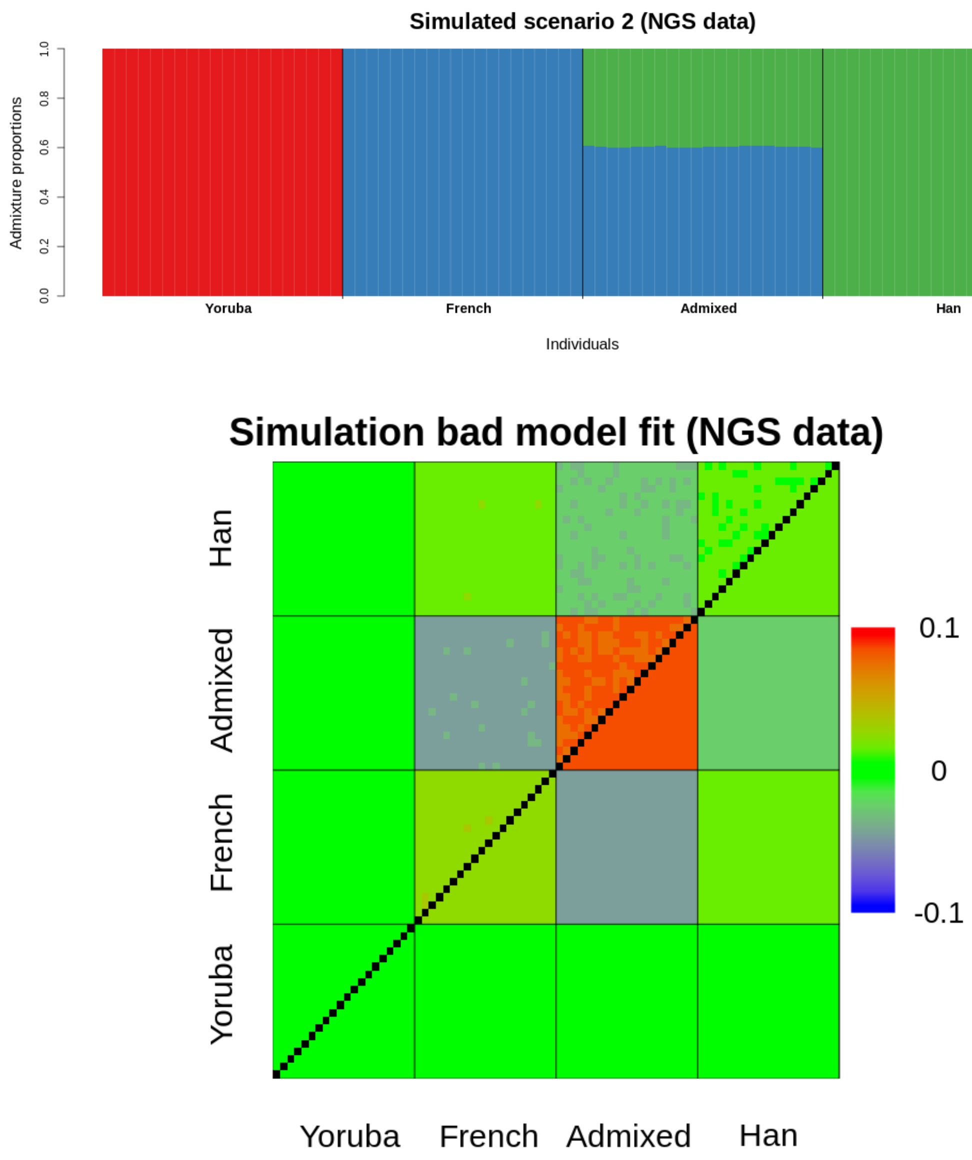


Figure S2: Admixture proportions inferred with NGSadmix assuming $K = 3$ from genotype likelihood data for simulated Scenario 2 (upper panel) and evaluation of model fit as correlation of residuals (bottom panel). Correlation values below or above the color scale are plotted as dark blue and dark red, respectively.

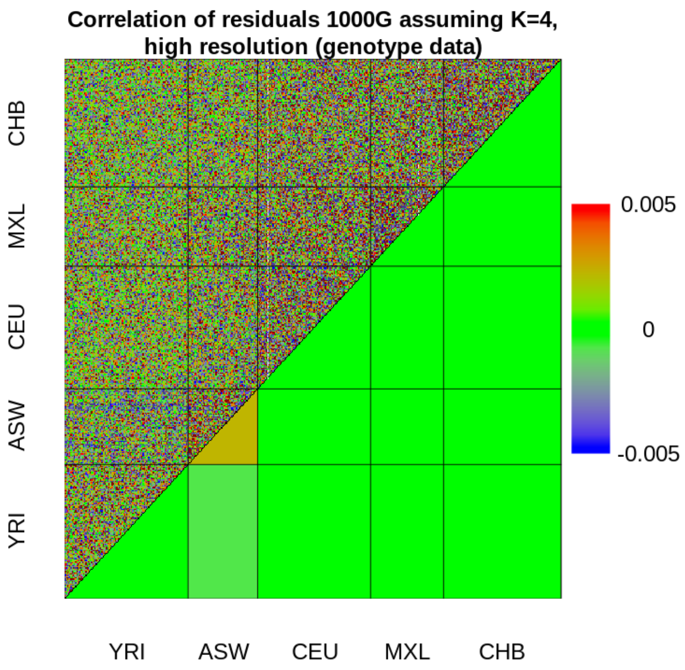


Figure S3: Evaluation of the inferred admixture proportions of the 1000G genotype data, assuming $K = 4$ and reducing the scale of the heatmap to highlight the presence of a positive correlation in the mean within ASW individuals. Correlation values below or above the color scale are plotted as dark blue and dark red, respectively.

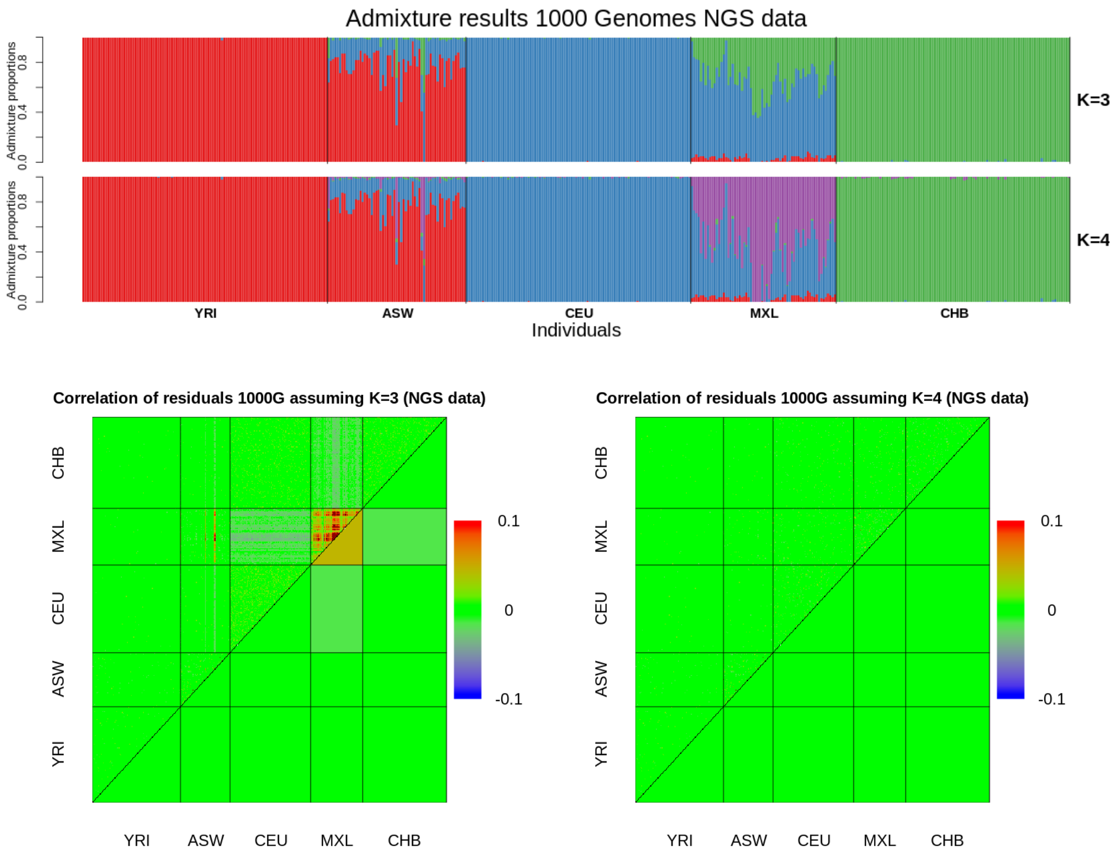


Figure S4: Inferred admixture proportions of the genotype likelihood data from the 1000G dataset, assuming $K = 3$ and $K = 4$ (upper panel). Evaluation of the estimated admixture proportions as correlation of residuals in each case, showing that four ancestral populations are needed to accurately model the data (bottom right panel, in contrast to bottom left panel). Correlation values below or above the color scale are plotted as dark blue and dark red, respectively.

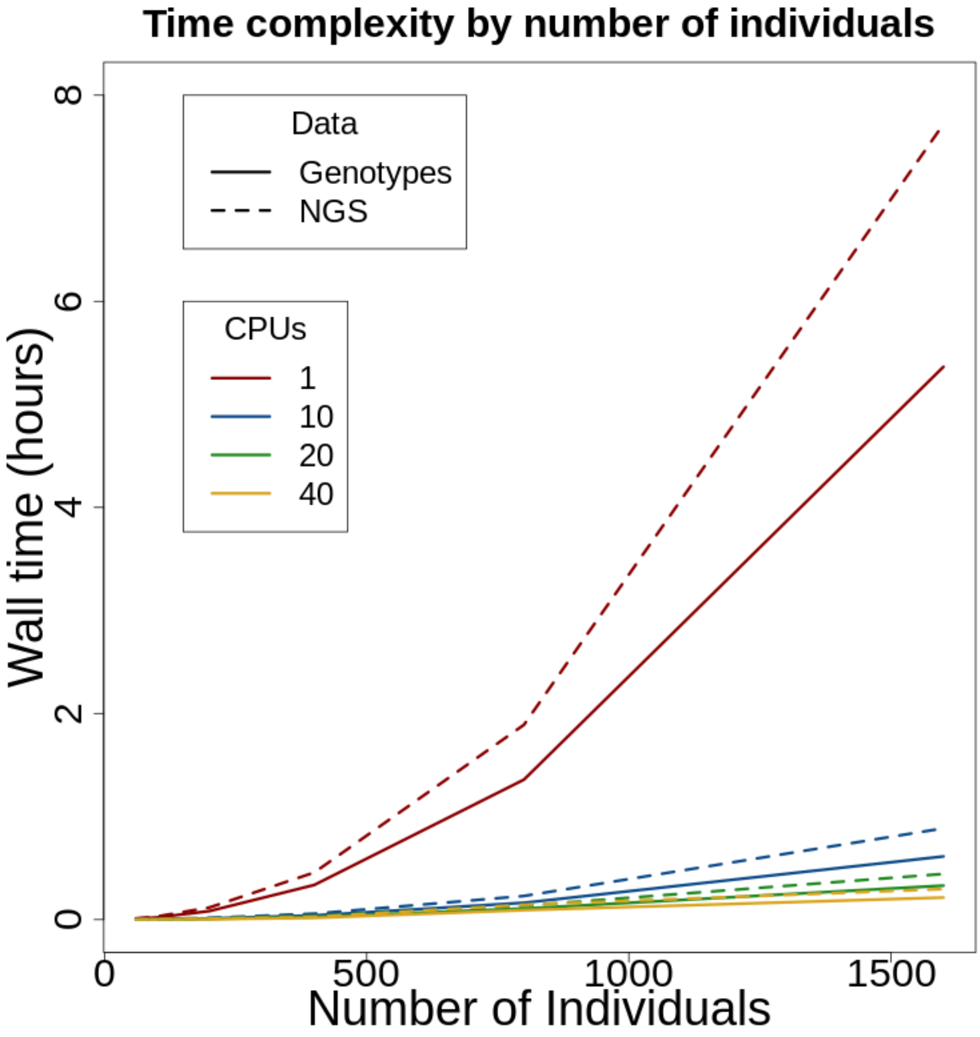


Figure S5: Total runtime of the genotype and NGS version of the program, with different number of individuals (N), and using 1, 10, 20 or 40 threads. In all cases the number of sites (M) is 100000 and the number of ancestral populations (K) is 6.
